# Fixed effect variance and the estimation of the heritability: Issues and solutions

**DOI:** 10.1101/159210

**Authors:** Pierre de Villemereuil, Michael B. Morrissey, Shinichi Nakagawa, Holger Schielzeth

## Abstract

Linear mixed effects models are frequently used for estimating quantitative genetic parameters, including the heritability, of traits of interest. Heritability is an important metric, because it acts as a filter that determines how efficiently phenotypic selection translates into evolutionary change. As a quantity of biological interest, it is important that the denominator, the phenotypic variance, actually reflects the amount of phenotypic variance in the relevant ecological stetting. The current practice of quantifying heritability from mixed effects models frequently deprives the heritability of variance explained by fixed effects (often leading to upward-bias) and it has been suggested to omit fixed effects when estimating heritabilities. We advocate an alternative option of fitting complex models incorporating all relevant effects, while including the variance explained by fixed effects into the estimation of heritabilities. The approach is easily implemented (an example is provided) and allows corrections for the estimation of heritability, such as the exclusion of variance arising from experimental design effects while still including all biologically relevant sources of variation. We explore the complications arising depending on the nature of the covariates included as fixed effects (e.g. biological or experimental origin, characteristics of biological covariates). Furthermore, we discuss fixed effects in non-linear and generalized linear models when fixed effects. In these cases, the variance parameters depend on the location of the intercept and hence on the scaling of the fixed effects. Integration over the biologically relevant range of fixed effects offers a preferred solution in those situations.

## Introduction

Additive genetic variance, phenotypic variance and their ratio, the heritability of a trait, are key parameters in evolutionary quantitative genetics, because they allow the assessment of whether a phenotypic trait can evolve through natural and artificial selection (Falconer and Mackay, 1996; Lynch and Walsh, 1998). The heritability, *h*^2^, of a trait corresponds to the fraction of the selection differential that can cause genetic change in the offspring generation. The heritability thus acts as a filter that determines how efficiently a population can respond to phenoytpic selection. Heritability is thus especially of interest to measure the adaptive potential of e.g. species threatened by global change (Hoffmann and Sgrò, 2011; Alberto et al., 2013), as well as to investigate fundamental issues in evolutionary biology (Mousseau and Roff, 1987; Merilä and Sheldon, 2000; Kruuk et al., 2000; Hadfield et al., 2006).

Mathematically, the heritability *h*^2^ of a trait is defined as the ratio of its additive genetic variance *V*_A_ to its total phenotypic variance *V*_P_:

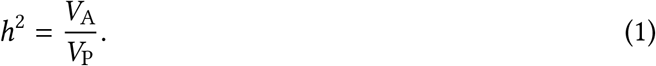

As a measure of biological interest, the heritability should be estimated with the ecologically relevant phenotypic variance in the denominator, just as *V*_A_ should be estimated accounting for various confounding effects (Wilson et al., 2010) and in the relevant environment, since genotype-by-environment interactions are common (Falconer, 1952; Kawecki and Ebert, 2004; Stinchcombe, 2014). The phenotypic variance *V*_P_ should ideally be quantified by random sampling from the base population in a biologically relevant setting. But studies are often designed, for good reasons, primarily for estimating the additive genetic variance *V*_A_ without bias and with the highest possible precision. Optimal sampling for the estimation of *V*_A_ can sometimes conflict between the precise estimation of the numerator and the denominator of Eq. 1. To cope with these design choices, as well as to model experimentally and naturally arising confounding effects, quantitative genetic models have to be as thorough as possible. This thoroughness inevitably leads to much complexity in the types and forms of effects included in the model, which in turns might render the computation of some parameters, especially *V*_P_, more difficult than usually appreciated.

The most popular methods for estimating quantitative genetic parameters make use of the linear mixed models (LMM) framework. In particular the so-called animal model (Thompson, 1976), a special case of a mixed effects model, is widely used in plant and animal breeding (Gianola and Rosa, 2015) and has been increasingly used in wild population studies over the past decade (Postma, 2014). One of the greatest advantages of mixed models is that they allow accounting for various confounding effects (Kruuk, 2004; Wilson et al., 2010). A LMM fitted to explain a phenotype y can contain both fixed and random effects, which is conventionally written as:

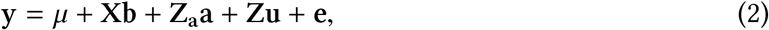

where y is the vector of phenotypes *y*, *μ* is the global intercept and **e** is a vector of residual errors. The **Xb** part stands for fixed effects (although not the intercept in the notation that we use here and in the following), whereas **Zu** refers to the random effects. Random effects, unlike fixed effects, are modelled as stemming from a Normal distribution with a mean of zero and a variance to be estimated from the data. Because of the quantitative genetic context discussed here, we isolate the random effect **Z_a_a** corresponding to the additive genetic value of the individuals from other random effect components. The matrices **X** and **Z** are referred to as the design and incidence matrices for the fixed and random effects, respectively. Especially, the **X** matrix contains the values of the co-factors included in the analysis. The vectors **b** and **u** contain the fitted fixed and potential random effect estimates, respectively.

When no fixed effects (apart from the intercept *μ*) are included in the analysis, the heritability is simply calculated as:

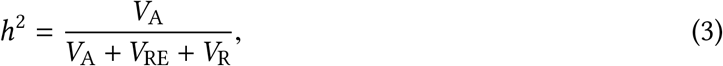

where *V*_A_ stands for the variance in additive genetic values **a**, *V*_RE_ for (the sum of) any additional random effect variance(s) and *V*_R_ for the residual variance. Since variance decomposition using LMM separates the phenotypic variance into additive components, Eq. 3 will generally give an unbiased estimate of Eq. 1. Fixed effects, however, can be problematic for multiple reasons and they are the focus of this paper.

Substantial progress has been made in highlighting issues pertaining to fixed effects in quantitive genetic inferences (Wilson et al., 2010; Wolak et al., 2015), generating solutions for mixed model analysis in general (Nakagawa and Schielzeth, 2013), and in data-scale quantitative genetic inference using generalised mixed models (de Villemereuil et al., 2016). Ideas in these works have resulted in substantial progress concerning the fitting and evolutionary quantitative genetic interpretation of mixed models with fixed effects. The purpose of this paper is to synthesise the ideas in these previous works so as to provide an accessible guidance to about what issues arise, and how to handle them, in a number of circumstances that are likely to occur in empirical evolutionary quantitative genetic studies.

## Heritability estimation in the presence of fixed effects

Fixed effects are often fitted with the intention to account for confounding effects and improve the goodness-of-fit of the models by accounting for complex patterns in the data. As illustrated in Fig. 1, the variance of the random effects, as well as the residual variance are estimated *around* the predicted values. Because of this, the sum of random effect and residual variances represents an underestimate of *V*_P_, as it does not reflect the total phenotypic variance of the trait, but rather the variance after the fixed effects have been accounted for (i.e. related to the red part in Fig. 1).

**Figure 1:**
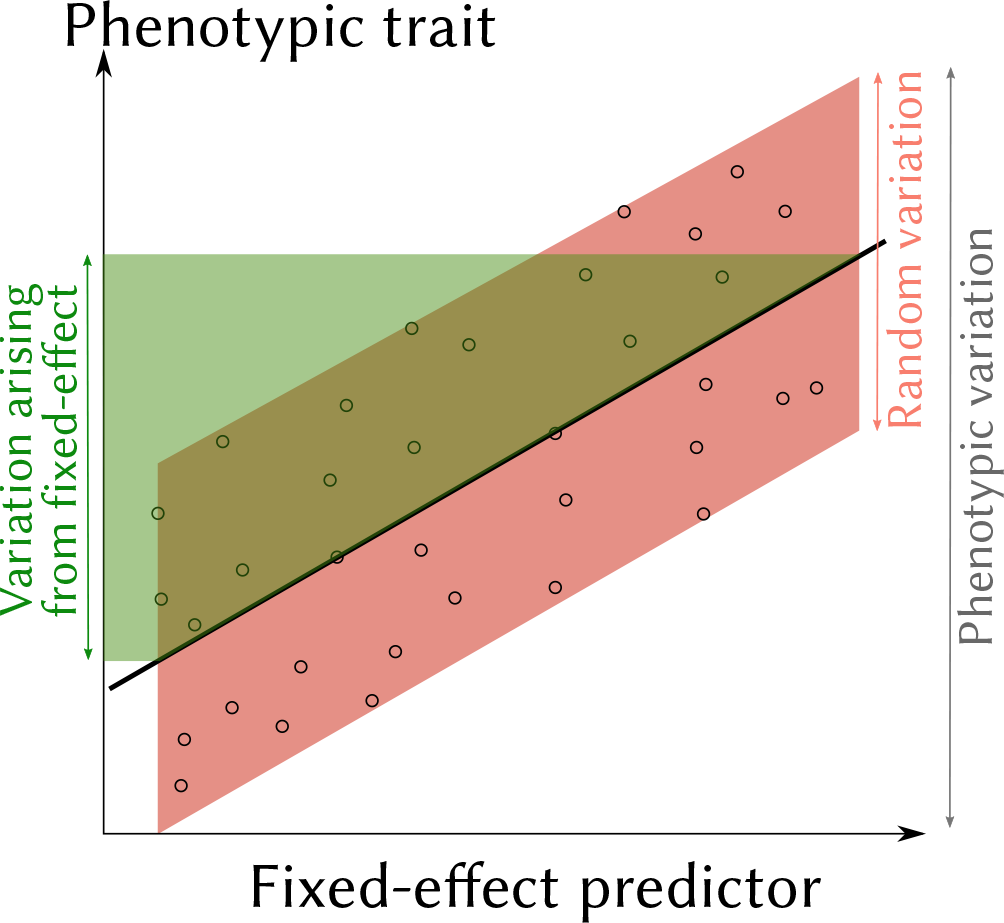
Schematic description of an analysis using a continuous fixed-effect predictor to model a phenotypic trait, possibly with random effects. The graph shows the relationship between the fixed-effect predictor and the phenotypic trait (individual data points in black circles, values predicted by the model as black thick line). The total phenotypic variation (black double-arrow on the right) is decomposed into the fraction explained by fixed-effect variation (i.e. the phenotypic variation “along” the predicted model, in green) on one hand, and random variation (i.e. variation from random effects and residual error arising “around” the predicted model, in red) on the other hand.

As a consequence, fixed effects affect the size of the phenotypic pie that is decomposed in different components, if the denominator is calculated as in Eq. 3. Wilson (2008) hence recommended particular care when fitting fixed effects in animal models and argued for a supplementary analysis without fixed effects. However, a cursory literature survey demonstrates that the use of fixed effects in quantitative genetics of wild populations is an almost universal practice (Table. 1). Note that the issues tackled here and by Wilson (2008) about reduction of the denominator variance when accounting for fixed effects also apply to the practice of two-step analyses by first fitting a linear model to account for confounding effects and then analysing the heritability of the residuals (Garland, 1988).

**Table 1:** Re-analysis of the Unicorn dataset from Wilson (2008), using models 1a, 1b and 1c from this reference. We computed *V*_F_ and provide *V*_P_ and *h*^2^ with or without accounting for this component. Discrepancies in values from *h*^2^ compared to Wilson (2008) are due a typological error in this reference (A.J. Wilson, personal communication).

| Fixed effect | $V_F$ | $V_A$ | $V_R$ | $V_P$ | | $h^2$ | |
| --- | --- | --- | --- | --- | --- | --- | --- |
| | | | | No $V_F$ | With $V_F$ | No $V_F$ | With $V_F$ |
| None | 0.00<br>[0.00-0.00] | 0.34<br>[0.08-0.60] | 3.14<br>[2.85-3.44] | 3.49<br>[3.24-3.71] | 3.49<br>[3.24-3.71] | 0.098<br>[0.025-0.17] | 0.098<br>[0.025-0.17] |
| Age + Sex | 2.49<br>[2.36-2.62] | 0.36<br>[0.27-0.47] | 0.65<br>[0.58-0.73] | 1.02<br>[0.94-1.08] | 3.51<br>[3.35-3.65] | 0.36<br>[0.28-0.45] | 0.10<br>[0.08-0.13] |
| Age + Sex +<br>Age:Sex | 2.56<br>[2.42-2.68] | 0.35<br>[0.26-0.46] | 0.60<br>[0.52-0.67] | 0.95<br>[0.88-1.03] | 3.51<br>[3.36-3.66] | 0.37<br>[0.28-0.46] | 0.1<br>[0.072-0.13] |

Since it will typically not be possible to get a benchmark for *V*_P_ from an independent dataset, we need solutions that allow a reconstruction of *V*_P_. A simple solution would be to replace the denominator *V*_A_ + *V*_RE_ + *V*_R_ by the phenotypic variance in the original dataset *V*_Po_, such that:

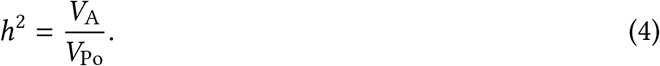

*V*_Po_ will however be affected by various aspects of the experimental design and may not be representative of the phenotypic variance in the base population (even if biases may be small in some cases of well-balanced experimental designs).

A more proper solution is to account for the amount of variance that has been transferred from the random components toward the fixed effects. In the context of computing the coefficient of determination (*R*^2^), Nakagawa and Schielzeth (2013) proposed to construct a fixed effect variance component as the variance of the linear predictor of the model 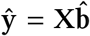 (where 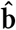 are the parameter estimates for the fixed effects). In other words, **ŷ** is the black thick line in Fig. 1 and its corresponding variance *V*_F_ (i.e. related to the green part in the figure) can be computed as:

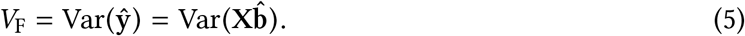

When including this variance component in the heritability calculation, the denominator is no more sensitive to the presence and number of fixed effects, because the variance transferred from random components to the fixed effects is now accounted for in the new component *V*_F_ (again, see Fig. 1 for a graphical illustration that *V*_P_ includes *V*_F_):

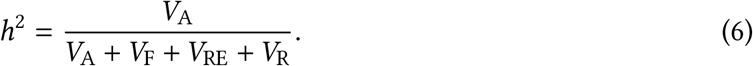

This is a straightforward calculation that can be done for any analysis, and using most software, since it needs only the values of the co-factors (i.e. the design matrices) and the parameter estimates. The former is an aspect of the sampling and/or experimental design and the latter is part of the output of any statistical software.

The inclusion of this *V*_F_ variance component in the computation of the total phenotypic variance *V*_P_ should be the norm when LMM are used to infer quantitative genetic parameters and when fixed effects are included in the analysis. It will be useful to provide estimates of this component in publications, in order to reflect how much variance was depleted because of the presence of fixed effects. The same kind of solution could be applied if the heritability was measured on the residuals of a regression (sometimes referred to as “corrected phenotypic values”): the variance of the regression model (*V*_F_, following the exact same definition as in Eq. 5) could be computed and included in *V*_P_, though a better practice anyway would be to run everything within a single LMM.

As an illustration, we re-analysed the Unicorn example data from Wilson (2008). This analysis (Table 1) shows that accounting for the *V*_F_ (note that values of *V*_F_ can be relatively high and certainly not negligible in general) component allow to recover the correct value for *V*_P_ and hence for *h*^2^, whichever the structure of the fixed effect component. Hence, because this practice would answer the concerns raised by Wilson (2008), we encourage researchers to include fixed effects in their analyses. A decision not to fit influential fixed effects, despite their beneficial effect on the goodness-of-fit or for accounting for confounding effects, would likely harm model fit, parameter estimation and the behaviour of the test statistics. Improving the fit of the model would most likely improve the precision of the estimates, which, for any particular dataset, would improve the precision of the heritability estimate (the point estimate would be more probably close to its true value and the confidence interval will be smaller). Furthermore, the inclusion of co-factors that account for nongenetic effects that are partly confounded with the additive genetic component *V*_A_ (e.g. common environment effects) are likely to reduce upward bias in the heritability estimate and will tend to result in lower, but more accurate point estimates of heritabilities.

## Removing the influence of experimental design on *V*_P_

In the context of estimating the phenotypic variance of a trait, fixed effects (as well as random effects) may be of two kinds. They can either reflect natural sources of variation that we are interested in, or variance arising from experimental and/or design effects. Since the latter category artificially inflates the variance in the data, we would, most of the time, like to exclude this source of variance from the heritability calculation. For example, if we want to study the amplitude of insect songs in the field, we might want to improve our model fit by including effects accounting for natural sources of variation, such as the age of the individual (if the amplitude is age-dependent) and effects accounting for sampling design, such as the distances between the animal and the microphone. Yet, in the computation of the phenotypic variance *V*_P_, we might want to include the biological variance arising from age, but not the experimental variance arising from the distance.

We have categorised fixed effects in the literature survey (Table 2) into sources of natural or experimental variation for illustration. Most of the fixed effects included in these analyses originate from natural variation (e.g. sex, year, age, area, litter size) and most likely should be included in *V*_P_. Others are of experimental origin either being an experimental treatment or of design origin (e.g. due to variation in the time of measurement) and should be excluded of *V*_P_. Of course, this separation between natural and experimental sources of variation can be quite difficult (e.g. year of sampling may represent error measurement or relevant natural variation depending on context). Furthermore it can sometimes be interesting to also exclude natural sources of variation. For example, “age” or “sex” could be excluded from the denominator to get heritabilities conditional on those factors. This would allow to perform evolutionary prediction for a particular age-class or sex.

**Table 2:** Fixed effects literature survey. This literature survey does not claim completeness, but should include the vast majority of heritability estimates in wild population using the animal model.

| Référence | Nb. effects | Fixed effects |  |
| --- | --- | --- | --- |
|  |  | Natural variation | Experimental variation |
| Réale et al. (1999) | 2 | Sex, Years | — |
| Kruuk et al. (2000) | 2 | Age, Area | — |
| Milner et al. (2000) | 8 | Age, Parasite burden, Birth Year, Year of measurement, Birth type, Coat color, Horn type | Catch date |
| Kruuk et al. (2001) | 1 | — | Brood size manipulation |
| Merilä et al. (2001) | 1 | — | Brood size manipulation |
| Kruuk et al. (2002) | 1 | Age | — |
| MacColl and Hatchwell (2003) | 6 | Year, Sex, Helper, Hatch date, Area, Attempt | — |
| Sheldon et al. (2003) | 1 | Year | — |
| Hadfield et al. (2006) | 5 | Sex, Year, Hatch date | Carotenoid treatment, Immune treatment |
| Thériault et al. (2007) | 4 | Age, Year | Day of capture |
| Nilsson et al. (2009) | 3 | Age, Dyad | Brood size manipulation |
| Morales et al. (2010) | 3 | Eggmass, Age | Treatment |
| Charmantier et al. (2011) | 2 | Sex, Natal colony | — |
| Doligez et al. (2011) | 2 | Sex, Age | — |
| Lane et al. (2011) | 2 | Age-class, Year | — |
| Reid et al. (2011b) | 1 | Year | — |
| Reid et al. (2011a) | 2 | Year, Age-class | — |
| Evans and Sheldon (2012) | 3 | Sex, Age | Measurement day |
| Béréños et al. (2014) | 3 | Sex, Litter size | Age at capture |
| Lane et al. (2015) | 4 | Age, Cone availability, Litter size, Year | — |

To exclude some particular factor(s), the predictor(s) (i.e. the respective columns in the design matrix) and the related inferred parameters can simply be left out of a new linear predictor 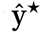 in the calculation of *V*_F_ such that:

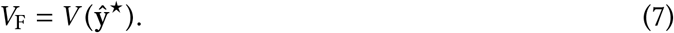

Note, however, that this computation is unfortunately not general and is based on the assumptions that the measured variance of the natural predictors is not “caused” (in the statistical sense of the term) by any of the experimental predictors. A more general solution relies on path analysis and the assumption of a causal pathway between variables (see Box 1).

#### Box 1: Using path analysis to obtain a partial *V*_F_

**Path analysis** Path analysis is a statistical analysis aiming at evaluating the *directed* influence of variables onto others. This directed relationship is referred to as *causality* (Wright, 1921). The direction of the relationship has a strong influence in our case, because it allows us to predict if the presence of one variable would inflate the variance of another.

**Three examples** In the figure below are three different examples using a phenotypic variable of interest ***P*** influenced by a biological variable ***B*** and an experimental variable ***E***. The parameters *b_XY_* stand for the coefficient of a model of the effect of *X* on *Y* (e.g. a slope). The parameters *σ_X_* is the exogenous standard-deviation of the variable *X*, i.e. its standard-deviation due to influences outside of the causal pathway (e.g. stochasticity, unmeasured variables and measurement error). The parameters *σ_XY_* is the exogenous covariance between *X* and *Y*, i.e. a undirected covariance arising from common influences outside of the causal pathway or due to physical/logical constraints (e.g. size and volume are physically covarying).

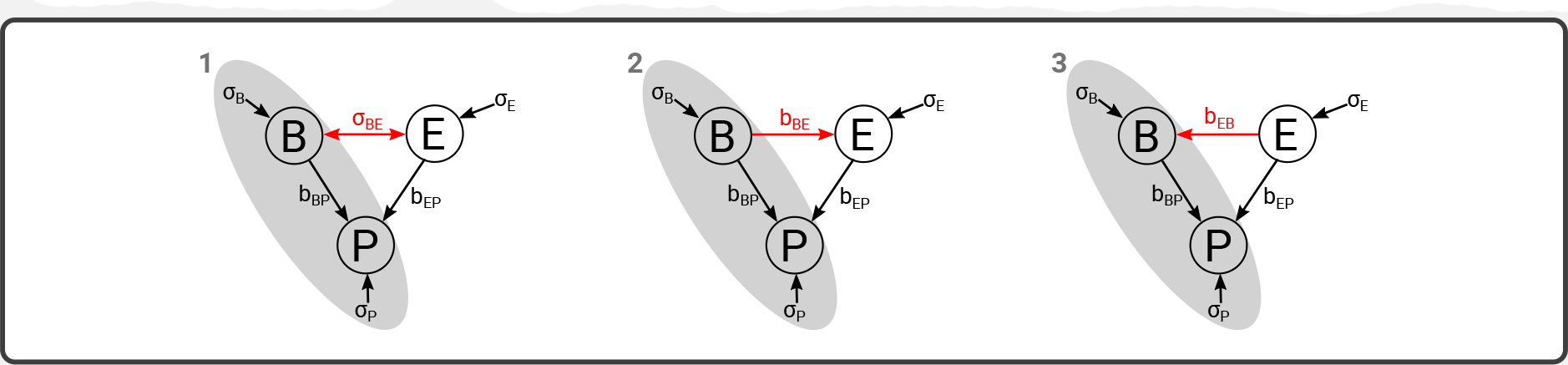

**General principle** In all cases, we are only interested in computing the variance arising from the grey area of the pathway (***B*** and ***P***), while excluding variance arising from ***E***. Excluding ***E*** from the graph means that we set its exogenous standard-deviation (*σ*_*E*_) and possible covariances (e.g. *σ*_*BE*_), as well as all the coefficients of its effect on any variable (e.g. *b*_*EP*_), to zero. Given that, the “fixed-effect variance” of ***P*** in this graph excluding ***E*** is simply the variance arising from the effect of *B*:

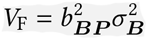

We will see that the difference between the three examples lies in the computation of *σ_B_*.

**Example 1** In this example, the variables ***B*** and ***E*** share an undirected covariance *σ_XY_*. In other words, we assume that a set of unmeasured variables have an effect on both ***B*** and ***E***, but not that a change in ***E*** will affect ***B***. In that case, the exogenous variance of ***B*** is merely its actual variance: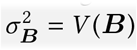.

**Example 2** In this example, the variable ***B*** has a direct effect on ***E*** (e.g. because an aspect of the species biology modulate the effect of the experimental treatment). In that case, changes of variance in ***B*** will affect the variance of ***E***, but this is not a problem for us since we want to exclude ***E***. Once again, the exogenous variance of ***B*** is merely its actual variance: 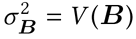.

**Example 3** In this example however, the variable ***E*** has a direct effect on the variable ***B*** (e.g. because the experimental treatment has an effect over different parts of the biological system). This means that, by experimentally introducing ***E*** into the biological system, we also experimentally increased the actual variance of ***B***. To compute the exogenous variance of ***B***, we need to remove this additional variance: 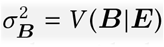. In other words, 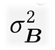 is here the residual variance of a model of the effect of ***E*** on ***B*** (e.g. the residual variance of the regression of ***E*** on ***B***).

In some rather special situations, it might be even advisable to replace the design matrix **X** by a modified design matrix **X'**, which implies using predicted values 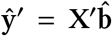 rather than fitted values in Eq. 5. For example, the insects that we are studying with respect to song amplitude might occur in distinct morphotypes (and these morphotypes might influence thermoregulation and thus song amplitude) that are not equally common. For statistical reasons it is advisable to oversample the rare morphs if we want to estimate the effect of morph on song amplitude. Such a sampling design will equalize morph frequencies in the sample and will thus tend to inflate *V*_F_ if calculated as 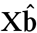 Since we argue that the denominator when calculating the heritability should represent the natural variation, we are better off with replacing **X** by **X'** that represents the natural frequencies of morphs in the population. Statistical requirements (balanced sampling) and biological realism (natural morph ratios) differ in this case and the calculation should account for this difference. Note that, while constructing **X'**, one should take into account potential correlations between co-factors. For example if the rare morph is preferentially present in warm environments and temperature is included in the model, then **X'** should reflect that correlation.

## Implicit assumptions about genetic covariances and fitting of genetic covariates

When fitting fixed effects in LMM, we are assuming stable residual and random-effect variances across the range of the covariates. If some covariates are of biological origin, we also implicitly assume a perfect genetic correlation along the range of those covariates. This assumption is frequently violated. In the special case of sex, for example, it has been shown that fitting sex as a fixed effect in a LMM leads to (downward) biased estimates unless the cross-sex genetic correlation is perfect (Wolak et al., 2015). But more generally, this applies to any factor or covariate that is added as a fixed effect.

A further consideration is whether fixed effects should cover only non-genetic sources of variation. Morphs in our example might be environmentally or genetically determined and it is usually advisable to model such discrete effects with potentially oligocausal control as fixed effects, no matter whether they are ultimately genetic or environmental in origin. With purely monogenic inheritance of morphs, morph phenotype is essentially a genetic marker for a (potential) quantitative trait locus (QTL) and thus represents the local heritability in linkage with the morph-determining locus (see e.g. Payne, 1918; Sax, 1923, for early QTL studies using Medelian phenotypes as markers), while the polygenic contribution of the background is captured by *V*_A_. Whether or not covariates cover genetic or non-genetic effects matters for the interpretation, since the estimate of *V*_A_ (and consequently *h*^2^) might represent the total *V*_A_ or the background *V*_A_ other than the local heritability at the QTL.

Some potential covariates might also be (heritable) polygenic traits themselves. In many cases, relationships between a focal trait and some other relevant trait are best handled with multi-response models (see Hadfield, 2010; Wolak et al., 2015), wherein the potential covariate is treated as a response along with the focal trait. Such a model estimates the genetic variances of, and genetic covariances (and others, e.g. residual) among the various traits treated as responses. This is not the case when the potential covariate is included as a fixed effect in the model: the fixed effect will explain the whole influence of the covariate on the focal trait but not explicitly distinguish between (nor differentially estimate) different sources of covariances. There are situations where it does make sense to include polygenic traits as fixed covariates, particularly when studying questions where causal effects of traits on one another are relevant. Further discussions of such scenarios are presented in Gianola and Sorensen (2004) and Morrissey (2014, 2015).

## Non-linear models and non-Gaussian traits

The influence of fixed effects might become more problematic when non-linearity is introduced in the model, in which case the approach proposed here will be inefficient. Such non-linearity arise obviously for non-linear mixed models (NLMMs), but also for generalised linear mixed models (GLMMs), through the non-linearity of their link functions. In GLMMs, there are indeed differences between the latent scale and the data scale (Morrissey et al., 2014): on the former, we assume linearity, normality and perform most of the inferences, whereas the latter is a non-linear transformation from the latent scale (e.g. through the link function). Hence, the above framework could be used on the linear, normally distributed, latent scale, but not with methods transforming estimates from the latent scale to the data scale like those reviewed in Nakagawa and Schielzeth (2010).

The non-linearity indeed breaks the assumption of independence between fixed effects and random effects, with the direct consequences that quantitative genetic parameters can no longer be computed without accounting for the whole distribution of fixed effects. Hence *V*_A_ and *V*_P_ on the data scale become complex functions of all the other parameters, rendering the computation of *V*_F_ essentially meaningless for this scale (see Fig. 2). De Villemereuil et al. (2016) showed that fixed effects must instead be integrated over to accurately compute quantitative genetic parameters using a GLMM. Integration over fixed effects is the solution to marginalize over the fixed effects and thereby accounts for their shape and distribution (for details see de Villemereuil et al., 2016). This approach assumes that the distribution of fixed effects in the sample is representative for the base population of interest. Otherwise, the design matrices might need to be adjusted accordingly as described above, or a distribution for the predictors must be assumed. The same logic applies when working with non-linear models or with data that was non-linearly transformed, unless we are specifically interested in the heritability of the transformed data.

**Figure 2:**
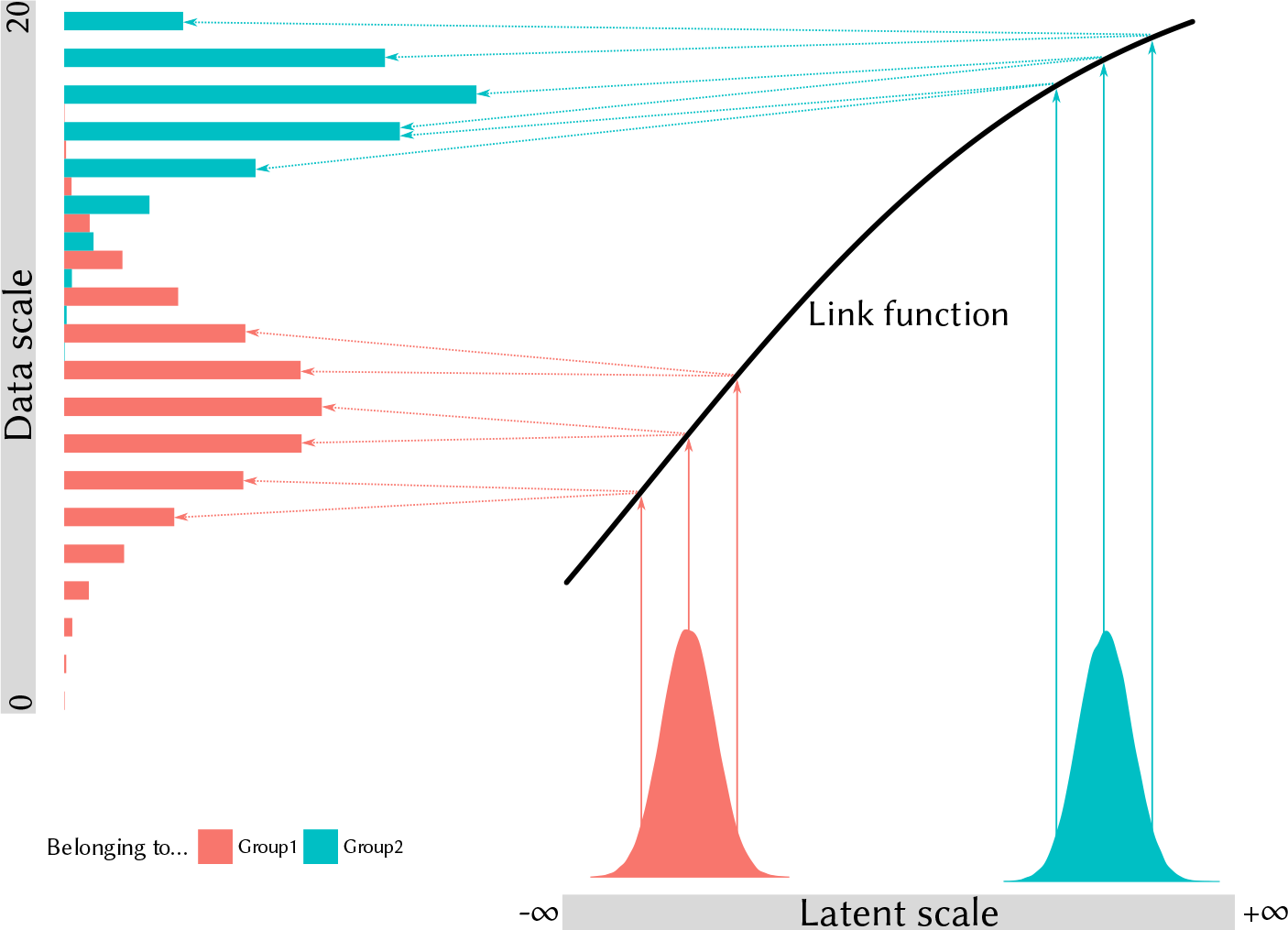
The figure illustrates the case of a binary trait with 20 observations per record that is modelled using a Binomial GLMM with logit link. Plain arrows illustrate deterministic relationships and dotted arrows stochastic relationships. On the latent scale, a fixed effect is assumed, accounting for the presence of two different groups (possibly males and females). The latent scale values for individuals of these two groups are clearly separated. According to the canonical assumptions of linear modelling, the groups only differ in their mean, not in their distribution or variance. Because the link function is not linear, initially equally apart values for each groups are more spread for Group 1 than for Group 2 (solid arrows). A further effect is the effect of the binomial, yielding more variance for medium probabilities (Group 1, dotted arrows) than for high probabilities (Group 2, dotted arrows). The end result is that on the data scale, the two groups no longer satisfy the assumptions on the latent scale: their variance are different (bigger for Group 1), the shapes of their distributions now differ (Group 2 is more skewed) and in this case the they two distributions even overlap. On the data scale, it is not possible to compute the variance arising from the fixed effect as simply the variance arising from differences in mean between the two groups.

It is thus important to stress that the approach suggested here can only be applied to phenotypic traits with a Normal distribution and analysed using linear mixed models (or if the analysis is based on the latent scale of a GLMM). However, the strategies presented here to remove the influence of experimental design still apply for GLMMs: experimental or sampling design effects can (and most likely should) be left out during the computation of the linear predictors and can be virtually “re-sampled” to account for biased sampling unrepresentative of natural populations.

## Conclusion & Perspectives

Wilson (2008) identified an issue when fixed effects are included a quantitative genetic model: the inclusion of fixed effects in the model has an influence on the computation of the phenotypic variance. Based on recent work from several sources, we provided guidelines to overcome this and other related issues, in the hope this will facilitate the use and interpretation of mixed models with fixed effects. We also discussed the complications arising from the diverse and complicated nature of covariates that can be fitted as fixed effects. We think that fixed effects are an opportunity to finely control confounding effects. Yet, when belonging to the phenotypic variance, they need to be included in the denominator of the heritability. In order to do so, we here promote the practice of accounting for the “fixed-effect” variance component *V*_F_ (see Nakagawa and Schielzeth, 2013), which includes the variance of all or selected fixed effects to be added in the denominator of the heritability calculation. We include an example of analysis using simulated data (see Supplementary Information) and the R package MCMCglmm (Hadfield, 2016) to show how these calculations can be implemented and how they can affect the output (*h*^2^ estimates going from 0.66 when not including *V*_F_ in the denominator to 0.15 when including it in our example).

This approach has several advantages. First, it allows to overcome Wilson (2008) legitimate reluctance of including fixed effects in the model. When including *V*_F_ in the denominator, there is no issue of “lost variance”. Second, since we are now able to include fixed effects, we have gained a finer control on confounding effects on the additive variance. It also requires some careful consideration of which fixed effects represent experimental design effects and which are biologically relevant. Third, it provides us with the choice of whether or not to include effects in *V*_F_, depending on whether or not we deem them part of the natural phenotypic variance of the studied population. Fourth, as argued above for the case of morphotypes in the context of song amplitude, the calculation of *V*_F_ can accommodate some discrepancies between the analysed data and the actual population.

Overall we advocate for the inclusion of fixed effects in linear mixed models to estimate heritabilities when *(i)* this improves the goodness-of-fit of the model and/or helps to account for confounding effects and *(ii)* a carefully computed *V*_F_ component is included in the calculation of the denominator of the heritability. While this is generally also true for non-linear models and GLMMs, any model that involves non-linearity in the response to fixed effects will require particular attention and likely integration over their biologically relevant range in order to marginalize the influence of fixed effects.

## Acknowledgements

We thank A.J. Wilson for sharing his Unicorn example data for us to analyse. HS was supported by an Emmy Noether fellowship from the German Research Foundation (DFG; SCHI 1188/1-1). SN is supported by a Future Fellowship, Australia (FT130100268). MBM is supported by a University Research Fellowship from the Royal Society (London).

